# Vaginal microbial dynamics and pathogen colonization in a humanized microbiota mouse model

**DOI:** 10.1101/2023.02.09.527909

**Authors:** Marlyd E. Mejia, Vicki Mercado-Evans, Jacob J. Zulk, Samantha Ottinger, Korinna Ruiz, Mallory B. Ballard, Robert A. Britton, Kathryn A. Patras

## Abstract

Vaginal microbiota composition is associated with differential risk of urogenital infection. Although vaginal *Lactobacillus* spp. are thought to confer protection through acidification, bacteriocin production, and immunomodulation, lack of an *in vivo* model system that closely resembles the human vaginal microbiota remains a prominent barrier to mechanistic discovery. We performed 16S rRNA amplicon sequencing of wildtype C57BL/6J mice, commonly used to study pathogen colonization, and found that the vaginal microbiome composition varies highly both within and between colonies from three distinct vivaria. Because of the strong influence of environmental exposure on vaginal microbiome composition, we assessed whether a humanized microbiota mouse (^HMb^mice) would model a more human-like vaginal microbiota. Similar to humans and conventional mice, ^HMb^mice vaginal microbiota clustered into five community state types (^h^mCST). Uniquely, ^HMb^mice vaginal communities were frequently dominated by Lactobacilli or *Enterobacteriaceae*. Compared to genetically-matched conventional mice, ^HMb^mice were less susceptible to uterine ascension by urogenital pathobionts group B *Streptococcus* (GBS) and *Prevotella bivia*, but no differences were observed with uropathogenic *E. coli*. Specifically, vaginal *Enterobacteriaceae* and *Lactobacillus* were associated with the absence of uterine GBS. Anti-GBS activity of ^HMb^mice vaginal *E. coli* and *L. murinus* isolates, representing *Enterobacteriaceae* and *Lactobacillus* respectively, were characterized *in vitro* and *in vivo*. Although *L. murinus* reduced GBS growth *in vitro*, vaginal pre-inoculation with ^HMb^mouse-derived *E. coli*, but not *L. murinus*, conferred protection against vaginal GBS burden. Overall, the ^HMb^mice are an improved model to elucidate the role of endogenous microbes in conferring protection against urogenital pathogens.

**IMPORTANCE:** An altered vaginal microbiota, typically with little to no levels of *Lactobacillus*, is associated with increased susceptibility to urogenital infections, although mechanisms driving this vulnerability are not fully understood. Despite known inhibitory properties of *Lactobacillus* against urogenital pathogens, clinical studies with *Lactobacillus* probiotics have shown mixed success. In this study, we characterize the impact of the vaginal microbiota on urogenital pathogen colonization using a humanized microbiota mouse model that more closely mimics the human vaginal microbiota. We found several vaginal bacterial taxa that correlated with reduced pathogen levels but showed discordant effects in pathogen inhibition between *in vitro* and *in vivo* assays. We propose that this humanized microbiota mouse platform is an improved model to describe the role of the vaginal microbiota in protection against urogenital pathogens. Furthermore, this model will be useful in testing efficacy of new probiotic strategies in the complex vaginal environment.

## INTRODUCTION

The vaginal microbiota is inextricably tied to women’s urogenital health. Perturbations to the vaginal microbiota are implicated in the risk of adverse outcomes including preterm birth, pelvic inflammatory disease, urinary tract infections, and sexually transmitted infections (1–9). Across geographic, ethnic, and social demographics, the composition of the vaginal microbiota in reproductive age women is largely comprised of *Lactobacillus* spp. Beneficial effects imparted by *Lactobacillus* spp. include lactic acid-mediated acidification of the vaginal environment, hydrogen peroxide and bacteriocin production, direct adherence to the vaginal epithelium, and immunomodulatory activity, all of which may hinder pathogen growth and colonization (10–16). While *Lactobacillus* dominance is canonically considered a hallmark of vaginal health, approximately 1 in 5 women have a more diverse vaginal microbiota characterized by a greater proportion of facultative or strictly anaerobic taxa such as *Bacteroides* and *Prevotella* (17, 18). Frequently, women with these more diverse, non-*Lactobacillus* dominant, communities are asymptomatic; however, women experiencing symptoms such as abnormal discharge and odor are often diagnosed with bacterial vaginosis (BV). These more diverse communities have gained a reputation as dysbiotic due to their association with vaginal symptoms and heightened risk of obstetric and gynecologic complications (18–22).

Although understanding of the human vaginal microbiota, its association with reproductive health and disease, and insight into specific protective mechanisms have expanded in recent years, many questions remain regarding the mechanistic roles of microbe-microbe interactions and their interplay with vaginal physiology and host immunity. Lack of an *in vivo* model system that closely resembles the human vaginal microbiota remains a prominent barrier to mechanistic discovery. While mouse models have served a seminal role in delineating host-microbe interactions in reproductive diseases, the murine vaginal microbiota has only recently been defined and is quite distinct from that of women; *Staphylococcus succinus and Enterococcus* spp. are the most common members in C57BL/6J mice (23, 24), *Enterobacteriaceae* and *Proteus* spp. are dominant in CD-1 mice (25), and *Streptococcus* spp. and *Proteus* spp. have been observed in FVB mice (26). Conventional mice are poorly colonized by human vaginal *Lactobacillus* spp. and require multiple, high-inoculum doses to observe *in vivo* effects (27–29). Several studies have evaluated the ability of human vaginal microbial communities to colonize mice with low or no endogenous microbiota, but stable colonization by these human communities was not achieved (30, 31). Not only is there a need to better understand the dynamic of the vaginal microbiota in conventional mice, especially as it relates to disease modeling, there is also a dire need for an animal model that better recapitulates the human vaginal microbial environment and provides translational relevance (32, 33).

Here, we evaluated the impact of environment on the vaginal microbiome in conventional C57BL/6J mice born and raised at three distinct vivaria. To circumvent challenges with transient colonization of human vaginal microbes, we assessed whether a mouse model that achieved stable colonization with human microbes, the humanized microbiota mouse (^HMb^mice) (34), would exhibit a more human-like vaginal microbiota. We investigated the impact of environment (vivarium room) and estrous on vaginal microbiome composition of ^HMb^mice and determined susceptibility to vaginal colonization by three human pathobionts. We found that the murine vaginal microbiota is malleable in composition and that the distinct ^HMb^mouse vaginal microbiota is capable of conferring protection against human pathobionts group B *Streptococcus* and *Prevotella bivia* compared to mice with conventional vaginal microbiota.

## RESULTS

### The vaginal microbiota differs between vivaria and demonstrates high intra-colony variability

Both human and mouse studies have shown that, despite host genetic selection for certain microbiome features (35), environmental factors play a dominant role in microbial composition of the gut (36–43); however, it is unknown whether the vaginal microbiota is likewise affected. To test this, vaginal swabs collected from conventional C57BL/6J mouse colonies bred at three different institutions [Baylor College of Medicine (BCM), Jackson Labs, and the University of California San Diego (UCSD)] were subjected to 16S v4 rRNA amplicon sequencing. Like prior studies, mice from Jackson Labs were primarily dominated by *Staphylococcus succinus* and *Enterococcus* spp. (23, 24), with occasional appearance of *Lactobacillus* spp., *Corynebacterium* spp., or *Acinetobacter* spp. dominant communities (**Fig. 1A**). Similarly, BCM mice displayed *S. succinus, Acinetobacter* spp. and *Pseudomonadaceae* dominant communities, with rare occurrence of *Lactobacillus* spp. dominance. In contrast, UCSD mice vaginal samples were primarily composed of *Pseudomonadaceae* or a combination of *Pseudomonadaceae* and *Pasteurellaceae*. Primary taxa driving differences between individual mice were *S. succinus, Enterococcus* spp., and *Pseudomonadaceae*, with mice from BCM and Jackson clustering more closely together compared to the UCSD cohort (**Fig. 1B**). Analysis of Composition of Microbiomes (ANCOM) identified *Pseudomonadaceae* and *Pasteurellaceae* (most abundant in UCSD mice) and *Comamonadaceae* (most abundant in BCM mice) as significantly different OTUs between vivaria.

**Figure 1.**
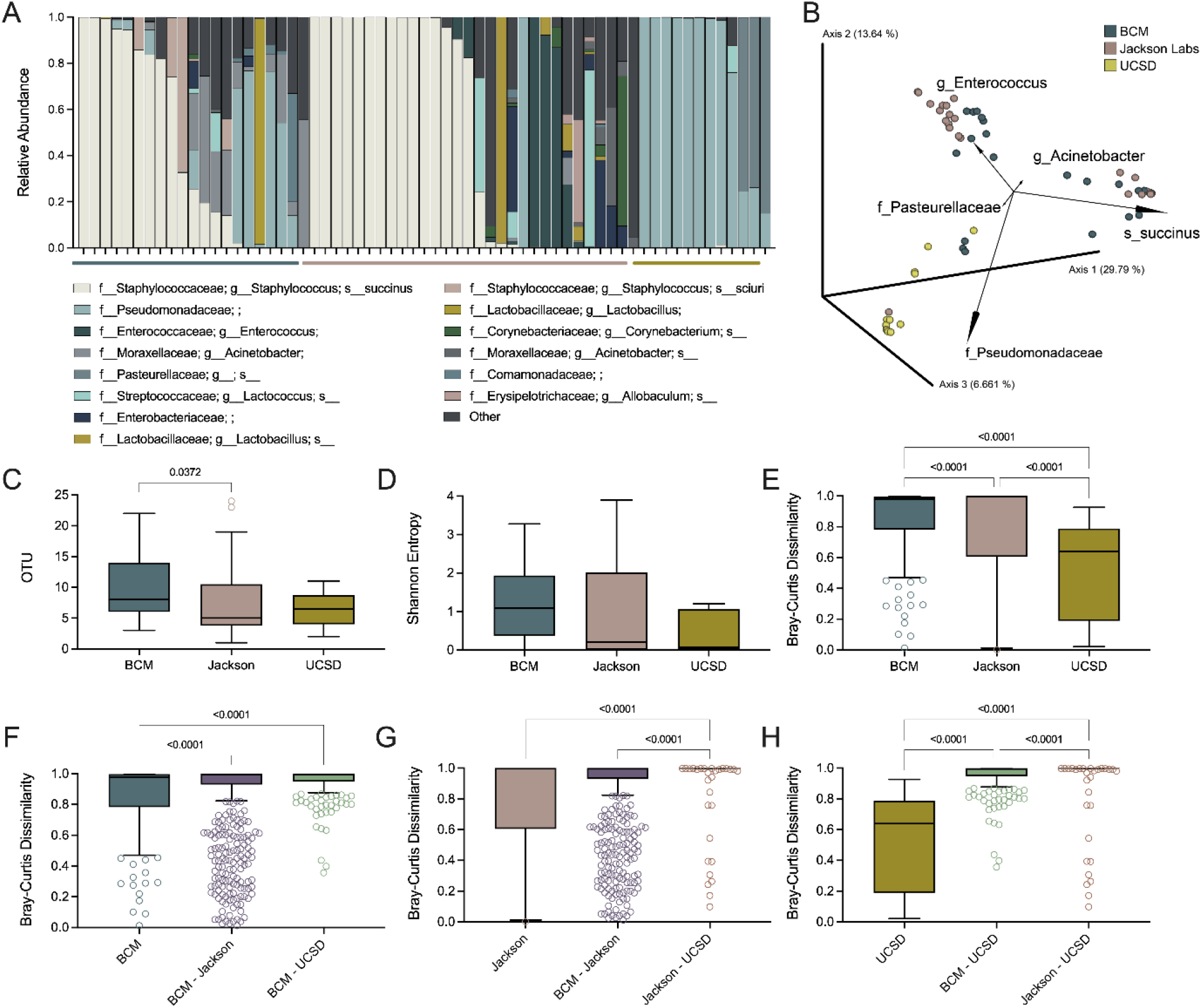
The microbial composition of the murine vaginal tract varies within colonies and between vivaria. Vaginal swabs from mouse colonies raised at BCM (*n* = 21), Jackson Lab (*n* = 30), and UCSD (*n* = 12) were collected and sequenced over the 16S rRNA V4 region. **(A)** Vaginal microbial compositions of mice at UCSD (yellow bar), Jackson Lab (pink bar), and BCM (blue bar). **(B)** PCoA plots of the Bray-Curtis distance matrix of individual mouse vaginal communities. **(C)** Observed OTUs in vaginal swab samples. **(D)** Shannon entropy of vaginal swab samples. **(E)** Dissimilarity between murine vaginal communities of mice within colonies (intra-site) at BCM, Jackson Lab, or UCSD. Intra-site variation compared to inter-site variation between BCM **(F)**, Jackson Labs **(G)**, and UCSD **(H)**. Dissimilarity plots were generated using PERMANOVA. Tukey’s boxplots are displayed. Each column **(A)** or symbol **(B)** represents a single mouse. Symbols and columns **(C-H)** represent pairwise Bray-Curtis distances. Data were analyzed by Kruskal-Wallis with a Dunn’s multiple comparison test and statistically significant *P* values are reported.

The number of observed OTUs was highest in the BCM colony, reaching statistical significance compared to the Jackson Labs colony (**Fig. 1C**), but no differences in Shannon entropy, a weighted alpha diversity metric, were observed (**Fig. 1D**). Intracolony variability was significantly different within each site, with UCSD having the most intra-colony similarity [Bray-Curtis distance (BC_med_) = 0.639] compared to BCM (BC_med_ = 0.978) and Jackson (BC_med_ = 0.998) (**Fig. 1E**). Inter-site dissimilarity was greater than that of intra-colony variability across institutional comparisons achieving statistical significance for most comparisons (**Fig. 1F-H**). Together, these data support that vaginal microbiota composition is heavily influenced by environment in the C57BL/6J genetic background, and thus manipulations of microbial or environmental exposures may alter the vaginal microbiota.

### ^HMb^mice have distinct vaginal communities compared to conventional mice and are enriched in *Lactobacillus-dominant* communities

To determine whether stable colonization of human-derived microbes in mice would alter the vaginal microbiota, we defined the vaginal microbiome of ^HMb^mice. ^HMb^mice, founded from germ-free WT C57BL/6J mice colonized with human fecal microbiota via oral gavage, display a more human-like gastrointestinal community compared to conventionally raised mice (34). The fecal microbiota of ^HMb^mice remains humanized over their lifespan and demonstrates generational stability of these communities among offspring (34). Vaginal swabs collected from multiple generations of ^HMb^mice over two years were subjected to 16S rRNA v4 amplicon sequencing. The first cohort of mice, twelve generations removed from founder mice, demonstrated *Lactobacillus* dominance (>70% relative abundance) in most samples (**Fig. 2A**). Subsequent cohorts had at least one mouse with *Lactobacillus* colonization (2.7-99% relative abundance), but the proportion of mice with *Lactobacillus* dominance decreased in cohorts 2-5 (**Fig. 2A**). Importantly, vaginal *Lactobacillus* in ^HMb^mice represent multiple species according to 16S v4 sequences. The most frequent *Lactobacillus* OTU sequences were identical to *L. fermentum, L. gasseri*, or murine-associated *L. murinus* by BLAST search (**Table S1**). Additional *Lactobacillus* OTU sequences mapped to *L. helveticus*, *L. zeae*, and *L. iners* which have been isolated from the human gastrointestinal and/or vaginal tracts (44–46). Other abundant taxa included *Enterobacteriaceae*, *S. succinus*, and *Enterococcus* spp. (**Fig. 2A, B**). When ordered by cohort, the greatest drivers of variation between mice were two distinct *Enterobacteriaceae* OTUs*, Acinetobacter* spp., and *S. succinus* (**Fig. 2B**). ^HMb^mice vaginal microbiota was notably distinct from the fecal microbiota in terms of composition and clustering by sample type on a PCoA of Bray-Curtis dissimilarities (**Fig. S1A-B**). Vaginal samples had decreased richness (median = 6.5 OTUs) compared to fecal pellets (median = 217 OTUs) (**Fig. S1C**) and lower alpha diversity determined by Shannon Entropy (**Fig. S1D**). Additionally, vaginal communities demonstrated greater variability between mice (BC_med_ = 0.998) compared to fecal communities (BC_med_= 0.463) (**Fig. S1E**).

**Figure 2.**
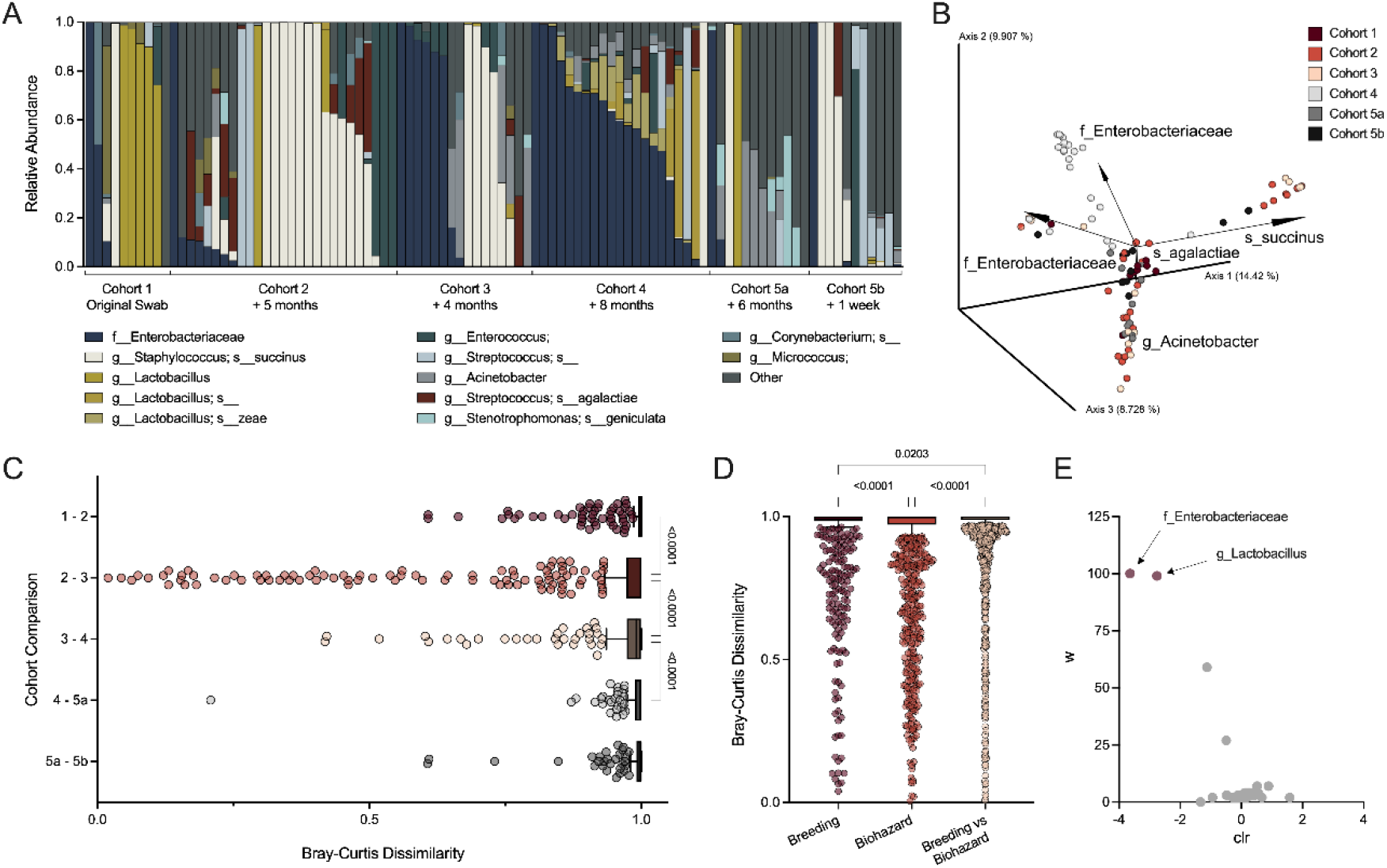
^HMb^mouse vaginal microbiota contains distinct taxa compared to conventional mice, is dynamic within the colony, and is sensitive to changes in vivarium. Vaginal swabs were collected from separate cohorts of ^HMb^mice over the course of two years. **(A)** Vaginal microbial compositions of distinct cohorts with the duration since previous sampling noted. Samples from Cohort 1 – Cohort 4 represent baseline vaginal swabs from unique mice (*n* = 10-27). Cohort 5 was swabbed at baseline (Cohort 5a) and sampled again one week after acclimation to the biohazard room (Cohort 5b). **(B)** Clustering of individual mouse cohorts according to Bray-Curtis Distances. **(C)** Dissimilarity between consecutive cohorts. **(D)** Dissimilarity between samples from mice swabbed in upon transfer from the breeding facility (*n* = 38) compared to those acclimated to and swabbed in the biohazard facility (*n* = 59). **(E)** ANCOM of taxa differentially abundant in the biohazard room (positive axis) and the breeding room (negative axis). Red points represent taxa that were found significant by ANCOM. Each column (A) or symbol (B) represents a unique mouse. Symbols **(C, D)** represent pairwise Bray-Curtis distances generated using PERMANOVA with Tukey’s boxplots displayed. Data for C and D were statistically analyzed by Kruskal-Wallis with Dunn’s multiple comparisons test and statistically significant *P* values are reported.

Compared to conventional mice, Bray-Curtis dissimilarity PERMANOVA measured high dissimilarity between ^HMb^mice and conventional mouse colonies for most permutations (**Fig. S2A**). PCoA visualization of ^HMb^mice overlaid with conventional mice showed separation of ^HMb^mice from UCSD and BCM mice, with some overlap of ^HMb^mice and Jackson mice that was partially driven by *Lactobacillus* spp. (**Fig. S2B**). Yet, as seen in conventional mice (**Fig. 1E**),^HMb^mice also demonstrated high variability across cohorts (**Fig. 2C**). The extent of dissimilarity between cohorts did not correspond with time between sampling periods. For example, dissimilarity between longitudinal samples from Cohort 5 taken only one week apart (Cohort 5a and 5b) was not different than that between two distinct sets of mice (Cohort 4 and Cohort 5a) taken six months apart (**Fig. 2C**). As part of the experimental design, animals were transferred from a colony (breeding) room to an experimental (biohazard) room in a detached vivarium, providing the opportunity to examine the vaginal composition shifted as the mice acclimated to the new vivarium. Although dissimilarity between mice within each room was high (BCavg = 0.937 in the breeding room and BCavg = 0.928 in the biohazard room), comparisons between rooms was even higher (BCavg = 0.963) suggesting divergent vaginal microbiota (**Fig. 2D**). Indeed, mice in the breeding room had higher abundances of *Lactobacillus* and *Enterobacteriaceae* than mice housed in the biohazard room for durations beyond one week (**Fig. 2E**).

### The vaginal microbiota in ^HMb^mice clusters into unique community states and minimally correlates with reproductive factors such as estrous stage

Vaginal microbial composition is implicated in birth outcomes in humans (47, 48) and fecundity in mice; reproductive success improves when germ-free mice become colonized with bacteria (49). To resolve whether ^HMb^mice have altered reproductive capacity, reproductive performance data were compared between conventional C57BL/6J mice at Jackson Labs (50), and genetically-matched conventional, germ-free, and ^HMb^mice colonies at BCM. The average age at first weaned litter in ^HMb^mice (14.4 weeks) was intermediate between conventional BCM mice (12.6 weeks) and germ-free BCM mice (20 weeks) (**Table 1**). Additionally, ^HMb^mice litter size (5.8 pups) was closest to that of conventional BCM mice (5.6 pups) compared to Jackson mice or BCM germ-free mice. ^HMb^mice were most similar to BCM germ-free mice in terms of total number of litters and gestational interval. Together, ^HMb^mice data fall within the range observed between conventional and germ-free mice implying that a humanized microbiota has minimal impact on reproductive performance. Reproductive differences between conventional BCM and Jackson mice were likely the result of colony management methods rather than biologic divergences.

**Table 1.** Reproductive parameters^a^ for C57BL/6J mice housed in different facilities and colonized by different microbial communities.

|  | **Conventional mice (Jackson Labs)<sup>b</sup>** | **Conventional mice (BCM)** | **Germ-free mice (BCM)** | **HMB mice (BCM)** | **HMB mice [range]** |
| --- | --- | --- | --- | --- | --- |
| Total # of litters per dam | 5.4 | 4.8 | 3.8 | 3.5 | [1 – 8] |
| Age of dam at first productive litter (weeks)<sup>c</sup> | 9.6 | 12.6 | 20.0 | 14.4 | [7.4 – 35.6] |
| Litter Size | 5.9 | 5.6 | 5.0 | 5.8 | [1 – 18] |
| Litters (total #/6 months) | 4.0 | 4.9 | 4.7 | 5.1 | [3 – 7] |
| Gestational interval (weeks) | Not specified | 4.6 | 6.8 | 6.8 | [2.9 – 19.3] |

<sup>a</sup>Mean values are reported.
<sup>b</sup>Data for Jackson Labs were taken from The Jackson Laboratory Handbook on Genetically Standardized Mice, 6<sup>th</sup> edition (50).
<sup>c</sup>First litters are typically lost despite being a productive mating. First successfully weaned litters are reported for BCM colonies.

The human vaginal microbiota displays modest fluctuations in composition and stability over the course of the menstrual cycle, including increased alpha diversity and decreased *Lactobacillus* relative abundance during menses (51–56). To determine if the ^HMb^mice vaginal microbiota is influenced by estrous stage, five ^HMb^mice were swabbed daily for one week. Relative abundances of taxa changed daily with each mouse displaying at least two different dominant taxa (>60% relative abundance) over the course of the week (**Fig. 3A**). Estrous stages were assigned by visualizing wet smears of vaginal samples as described previously (24) (**Fig. 3B**). To determine if estrous stage influenced vaginal microbiota composition, pairwise Bray-Curtis distances were calculated for vaginal samples (*n*= 34 mice, 1-7 samples/mouse) and grouped by the corresponding estrous stage at the time of sample collection (**Fig. 3C**). Although some clustering of similar communities were observed, these clusters were not associated with a particular estrous stage (**Fig. 3C**). Vaginal communities within each stage showed high dissimilarity, ranging from diestrus (BC_med_ = 0.93) to metestrus (BC_med_ = 0.993) (**Fig. 3D**). While richness did not differ between stages (**Fig. 3E**), alpha diversity was higher for mice in proestrus (Shannon_med_ = 2.13) and diestrus (Shannon_med_ = 2.08) compared to estrus (Shannon_med_ = 0.44) (**Fig. 3F**).

**Figure 3.**
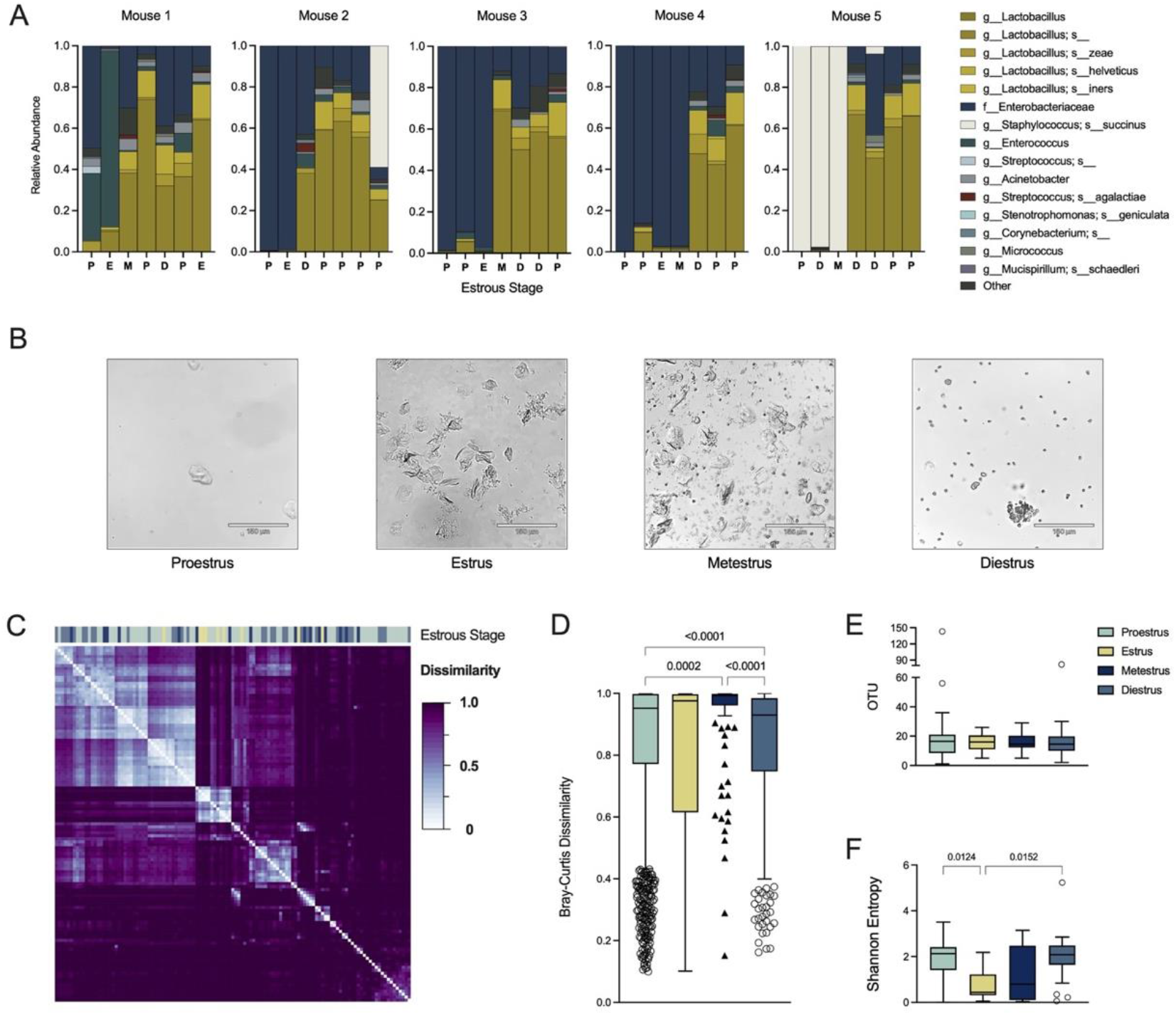
Vaginal microbiota dynamics over different estrous stages in ^HMb^mice. **(A)** Vaginal microbial compositions of five individual ^HMb^mice swabbed daily over the course of a week. The initials of estrous stage assignment at the time of sampling are denoted below the sample (P = proestrus, E = estrus, M = metestrus, D = diestrus). Only Mouse 1 and Mouse 2 were co-housed. **(B)** Representative microscopic images of vaginal wet smears collected at each stage of the estrus cycle. Wet smears from vaginal swab samples were visualized at 10X on a brightfield microscope. **(C)** Bray-Curtis distance matrix of vaginal swab samples colored by estrous stage (*n* = 119). Samples include 1-7 swabs per mouse (*n* = 34 mice). **(D)** Bray-Curtis dissimilarity of microbial compositions between samples categorized in the same estrous stage. **(E)** Observed OTUs and **(F)** Shannon diversity of samples grouped by estrous stage. Each column **(A)** represents a sample from Day 1 through Day 7. Symbols **(D)** represent pairwise Bray-Curtis distances. Open circles **(E, F)** represent individual mice outside Tukey’s whiskers. Data were statistically analyzed by Kruskal-Wallis with Dunn’s multiple comparisons test and statistically significant *P* values are reported.

Vaginal samples across cohorts were hierarchically clustered into murine community state types using Ward’s linkage of Euclidean distances as done previously (23, 24). Because the dominant taxa differed from conventional mice, we designated ^HMb^mice profiles as “humanized mCST” (^h^mCST). Two communities resembled mCSTs of conventional mice; ^h^mCST II (*Staphylococcus succinus*-dominant) and ^h^mCST IV (heterogenous taxa with an even composition). However, ^h^mCST I was *Lactobacillus* dominant, ^h^mCST III was *Enterobacteriaceae*-dominant, and ^h^mCST V contained either *Enterococcus, Streptococcus*, or *Lactobacillus-dominant* communities (**Fig. 4A**). Proportions of each ^h^mCST were not significantly different between estrous stages (**Fig. 4B**). ANCOM was performed between combined proestrus and estrus (increased estrogen) stages and combined metestrus and diestrus (decreased estrogen) stages or by the transition stages diestrus and proestrus versus estrus and metestrus; neither comparison revealed significantly different taxa between stages. When plotted by Bray-Curtis distances, samples did not separate into discrete clusters based on estrous stage but did cluster by ^h^mCST which is driven largely by taxonomy (**Fig. 4C**). Notably, there were two separate *Enterobacteriaceae-*driven clusters (**Fig. 4C**).

**Figure 4.**
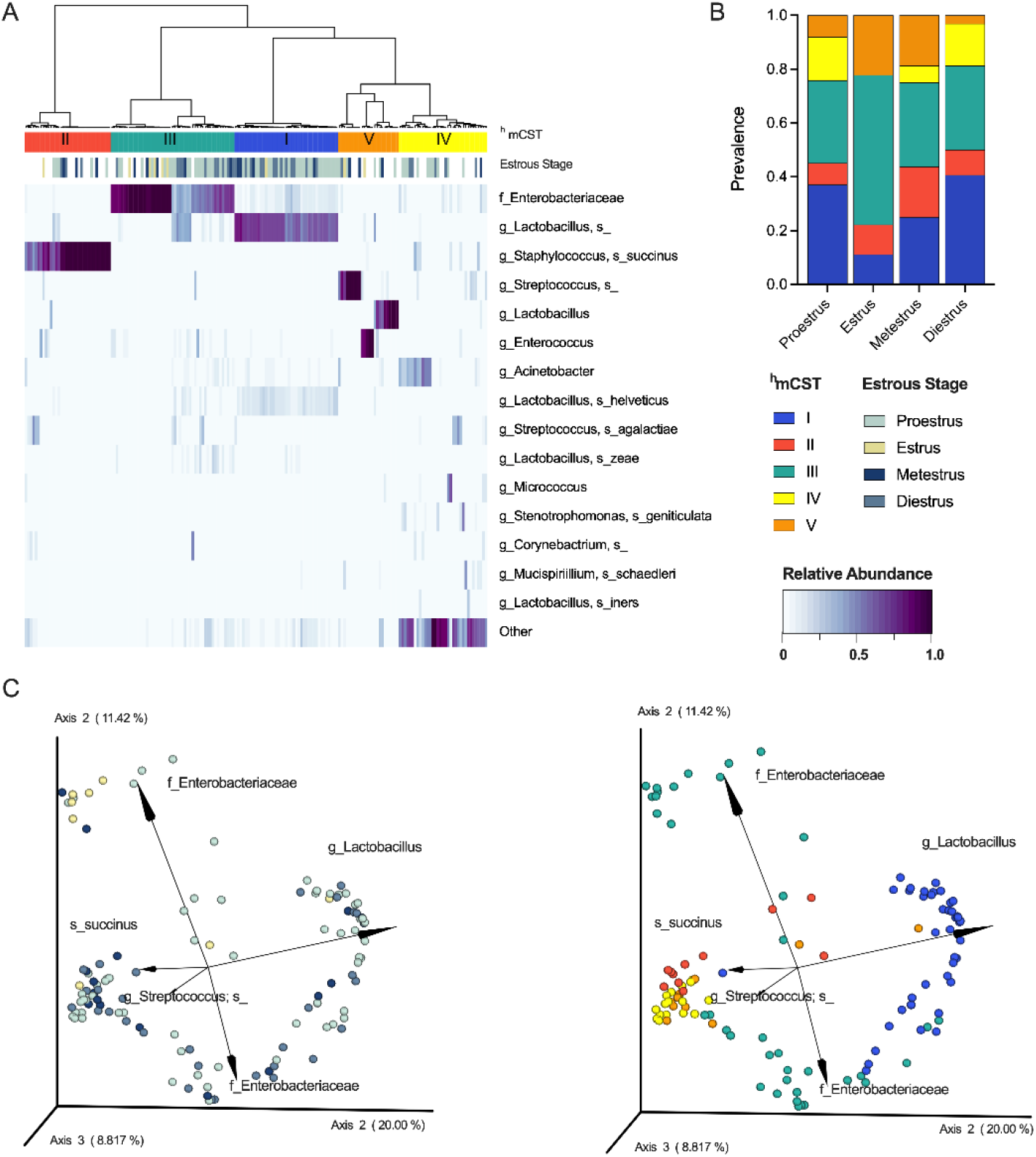
^HMb^mice exhibit distinct community state types that are not associated with specific estrous stages. Vaginal swabs were subjected to paired estrous staging and 16S rRNA sequencing. **(A)** Community state type categorization for ^HMb^mice (*n* = 183 samples) hierarchically clustered into humanized murine CST (upper bar) and its associated estrous stage (lower bar). ^HMb^mice include mice (*n* = 34) from **Fig. 3C** and additional mice (*n* = 64) that were sampled for 16S sequencing but were not staged. **(B)** Prevalence of ^h^mCSTs in each estrous stage. **(C)** Clustering of ^HMb^mice (*n* = 34) vaginal communities sampled over the course of a week according to estrous stage (left panel) and ^h^mCST (right panel). Each symbol **(C)** represents a unique mouse. Data in (B) were analyzed by Chi Square analysis of fractions. No values were statistically significant.

### ^HMb^mice exhibit decreased uterine ascension of group B *Streptococcus* compared to conventional mice

To determine whether the distinct ^HMb^mice vaginal microbiota impacted colonization by potential pathogens, we used an established colonization model of the neonatal pathogen group B *Streptococcus* (GBS) (57). GBS asymptomatically colonizes the maternal vaginal tract, but perinatal exposure during pregnancy or labor and delivery can cause severe disease including stillbirth or neonatal sepsis (58). Conventional C57BL/6J mice and ^HMb^mice were vaginally inoculated with 10^7^ CFU of GBS and swabbed daily over seven days (**Fig. 5A**). At early time points, ^HMb^mice had similar or higher GBS colonization compared to conventional mice. At later time points, however, some ^HMb^mice cleared GBS below the limit of detection resulting in significantly lower vaginal GBS burdens than conventional mice at Day 7 (**Fig. 5B**). To assess GBS ascension, reproductive tract tissues were harvested at Day 3 and Day 7. Uterine GBS burdens were significantly lower in ^HMb^mice compared to conventional mice at both time points, while vaginal and cervical GBS burdens were not different between groups (**Fig. 5C-D**).

**Figure 5.**
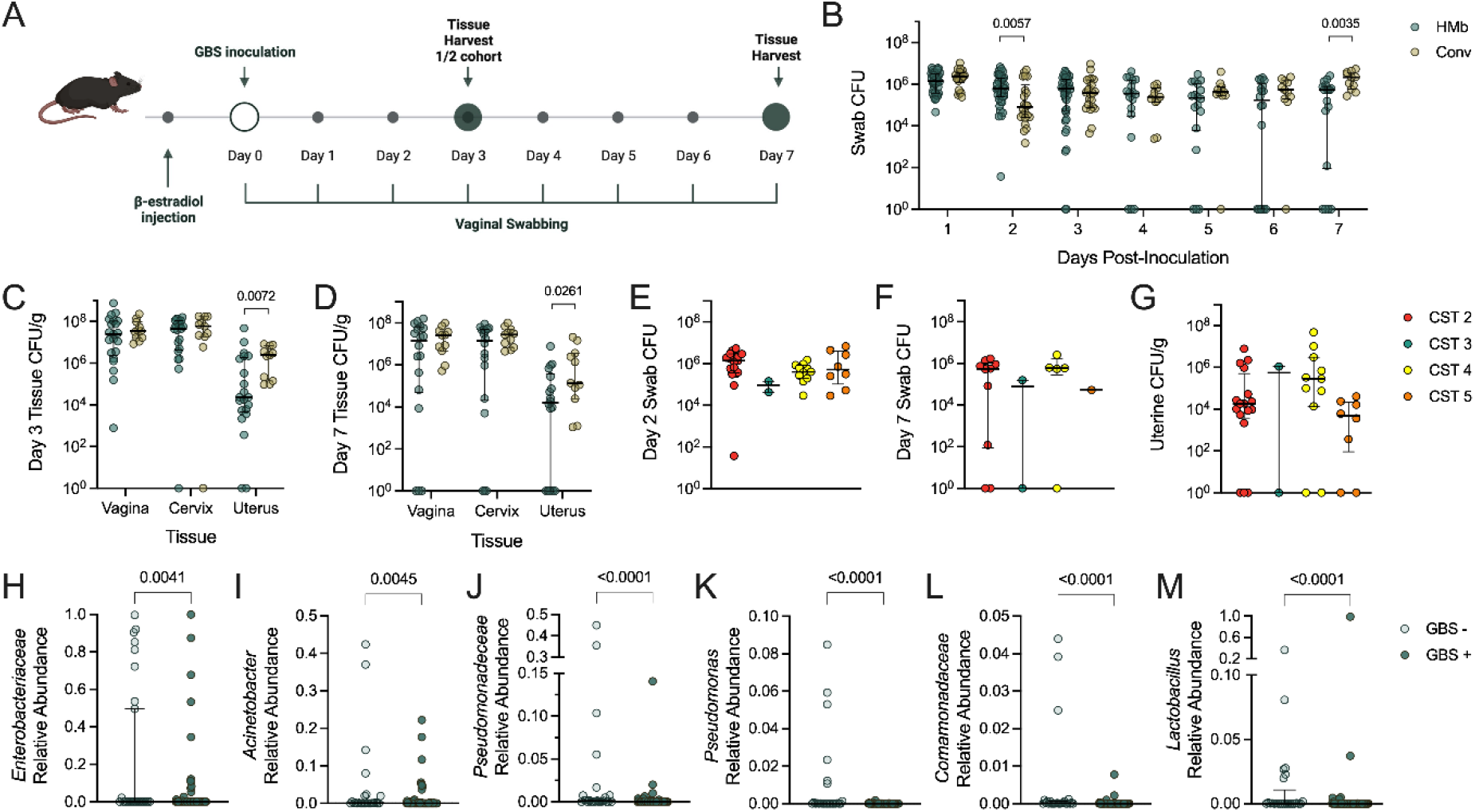
Increased vaginal clearance of GBS and restriction of uterine ascension in ^HMb^mice is not due to ^h^mCST but may be attributed to individual taxa. **(A)** ^HMb^mice (Hmb) and conventional (Conv) mice were vaginally inoculated with 10^7^ CFU of GBS (*n* = 10-27). **(B)** GBS CFU recovered from daily vaginal swabs. Vaginal, cervical, and uterine GBS tissue burdens were collected at **(C)** Day 3 and **(D)** Day 7 post-inoculation. GBS CFU counts from **(E)** Day 2 swabs, swabs, and **(G)** Day 3 and 7 uterine tissues delineated by ^h^mCST assignment of respective mice on Day 0 prior to GBS inoculation. Relative abundances of vaginal **(H)** *Enterobacteriaceae*, **(I)** *Acinetobacter*, **(J)** *Pseudomonadaceae*, **(K)** *Pseudomonas*, **(L)** *Comamonadaceae*, and **(M)** *Lactobacillus* across all vaginal swabs in mice group into detectable uterine GBS (GBS+) or no detectable uterine GBS (GBS-) at the time of tissue collection. Symbols represent individual mice. Data were statistically analyzed by Mann-Whitney test (A-C, G-L) or Kruskal-Wallis with Dunn’s multiple comparison test (D-F) and statistically significant *P* values are reported.

To determine if vaginal microbial profiles correlated with GBS burdens, ^HMb^mice GBS burdens were replotted according to the ^h^mCST assigned at Day 0 immediately prior to GBS inoculation. GBS vaginal burdens were not significantly different across ^h^mCST groups at Day 2 nor Day 7 (**Fig. 5E-F**). Furthermore, no significant differences in GBS uterine burdens from combined Day 3 and 7 samples were detected between ^h^mCSTs (**Fig. 5G**). To determine whether specific vaginal taxa were associated with GBS uterine ascension, ^HMb^mice were binned into two categories across both time points: those with no detectable GBS uterine CFU (GBS-) or those with detectable GBS uterine CFU (GBS+). Corresponding vaginal swab 16S sequences from all timepoints were then probed for differentially abundant taxa by ANCOM. Mice with no detectable uterine GBS exhibited an enrichment of *Enterobacteriaceae*, *Acinetobacter*, *Pseudomonadaceae*, *Pseudomonas*, *Comamonadaceae*, or *Lactobacillus* (**Fig. 5H-M**).

### *Lactobacillus murinus* and *E. coli* display discordant phenotypes towards GBS in competition assays *in vitro* and GBS vaginal colonization *in vivo*

To gain insight into mechanisms of differentially abundant taxa between groups, bacterial isolates were collected from mice with ^h^mCST I and ^h^mCST III communities as described in Methods. Two isolates, identified as *L. murinus* and *E. coli* by full-length 16S sequencing, were each cultured in MRS broth in competition with GBS at two timepoints across five different starting ratios. Minimal differences in competitive index were observed between *L. murinus* and GBS, with GBS displaying a significantly increased advantage at the 1:2 and 1:10 ratios at 3 h (*P* = 0.0102 and 0.0002 respectively) which was retained at the 18 h timepoint in the 1:10 condition (*P* = 0.0165) (**Fig. 6A**). To determine how coculture impacted growth of each organism, viable CFU of each organism in coculture was compared to CFU recovered from monoculture. *L. murinus*was minimally impacted (**Fig. 6B**), however, GBS growth at 18 h was impaired in the presence at *L. murinus* at all but the highest GBS starting inoculum (1:10 *L. murinus* to GBS) (**Fig. 6C**). Conversely, GBS demonstrated a strong competitive advantage in coculture with *E. coli*, which was significant in the 1:2 and 1:10 conditions at 3 h (*P* = 0.001 and <0.0001 respectively) and in all five ratios at the 18 h timepoint (*P* ≤ 0.0286) (**Fig. 6D**). Again, growth of each organism in coculture was compared to growth in monoculture. No differences were observed at the 3 h timepoint (**Fig. 6E**). At 18 h, *E. coli* growth was significantly impaired in the presence of GBS in all conditions (**Fig. 6F**). Raw viable CFU values for each organism are provided in **Fig. S3**.

**Figure 6.**
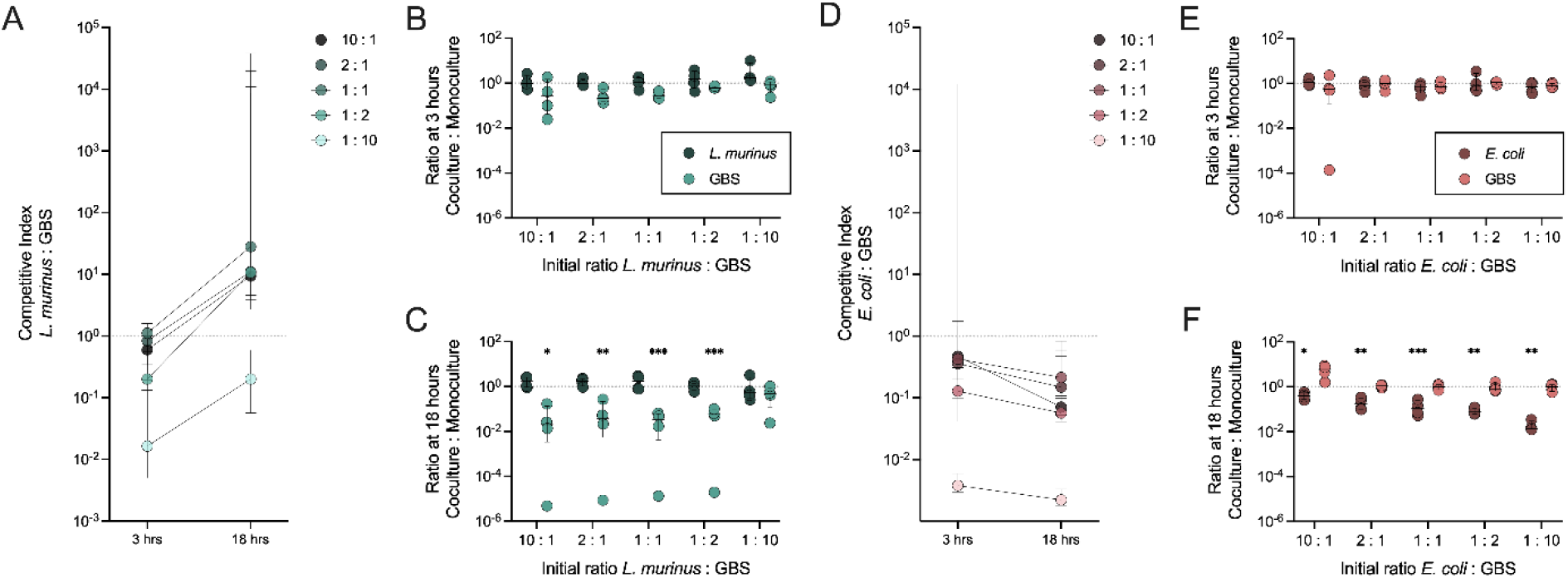
*In vitro* competition assays demonstrate GBS inhibition by *L. murinus* but not *E. coli*. *In vitro* competition assays were performed between GBS and**(A-C)** *L murinus* or **(D-F)** *E. coli*.**(A, D)** Competitive index (CI) was calculated as the CFU ratio of the treatment microbe over GBS, normalized to the initial ratio of the inoculum. The rate of bacterial growth (viable CFU) in coculture compared to monoculture controls was calculated at timepoints **(B, E)** 3 hours and **(C, F)** 18 hours. Symbols represent independent experimental replicates. Data were statistically analyzed by one sample t test with a theoretical mean of 1.0 (A, D) and two-way ANOVA with Šídák’s multiple comparisons test for coculture deviation from growth in monoculture (B-C, E-F); \**P*< 0.05; \*\**P* < 0.005; \*\*\**P* < 0.0005. Raw viable CFU values are reported in **Fig. S3**.

To validate ANCOM findings and determine whether pre-existing vaginal taxa could confer protection against GBS, *L. murinus* or *E. coli* were separately vaginally inoculated into ^HMb^mice prior to GBS challenge (**Fig. 7A**). Despite decreased growth of GBS in the presence of *L. murinus in vitro*, GBS vaginal colonization and dissemination into the upper reproductive tract was unaffected *in vivo* (**Fig. 7B-C**). Prophylactic inoculation with *E. coli* reduced GBS vaginal burden on Day 1 (2-log reduction) and Day 2 (3-log reduction) (**Fig. 7D**) but did not influence tissue burdens at Day 7 (**Fig. 7E**).

**Figure 7.**
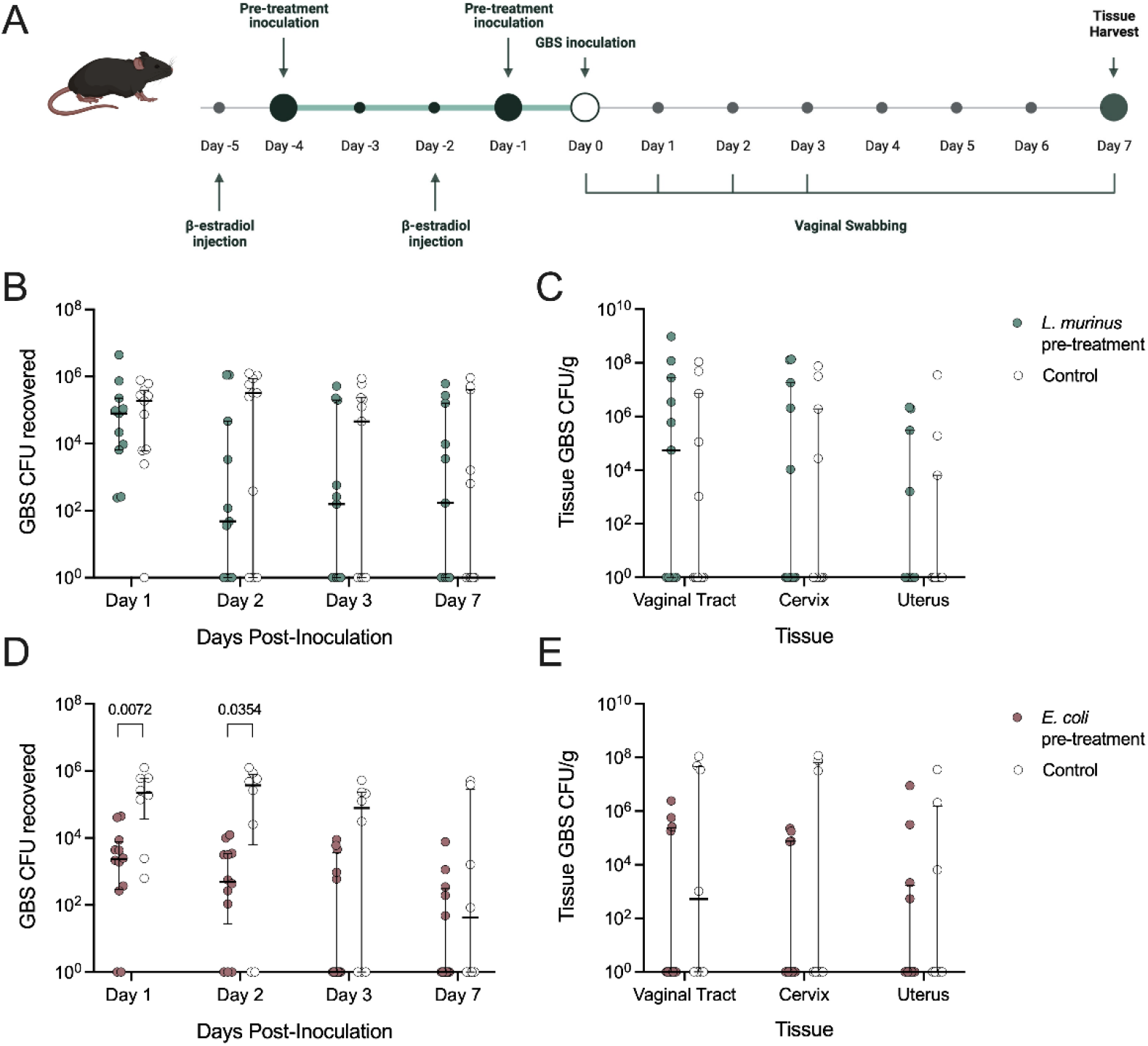
Pretreatment with *E. coli*, not *L. murinus*, reduces GBS vaginal colonization *in vivo*. **(A)** ^HMb^mice were vaginally inoculated with 10^7^ CFU of *L. murinus*,10^7^ CFU of *E. coli*, or vehicle control twice prior to challenge with 10^7^ CFU of GBS. Recovered GBS CFU from **(B)** vaginal swabs and **(C)** Day 7 vaginal, cervical, and uterine tissue of mice pre-inoculated with *L. murinus*. Recovered GBS CFU from **(D)** vaginal swabs and **(E)** Day 7 vaginal, cervical, and uterine tissue of mice pre-inoculated with *E. coli*. Symbols represent individual mice. Data were statistically analyzed by Mann-Whitney and significant *P* values are reported.

### ^HMb^mice exhibit decreased cervical and uterine ascension of *Prevotella bivia*, but not uropathogenic *E. coli*, compared to conventional mice

To determine whether ^HMb^mice were protected against multiple vaginal pathogens or selectively resistant to GBS, we challenged ^HMb^mice with two additional vaginal pathobionts associated with vaginal dysbiosis: *Prevotella bivia*, which is increased in women diagnosed with bacterial vaginosis (59), and uropathogenic *E. coli* (UPEC), a causative agent for urinary tract infection, which can establish vaginal reservoirs (60) or cause aerobic vaginitis (61). Unlike GBS, there was no difference in vaginal swab *P.bivia*burdens between conventional or ^HMb^mice at any timepoint (**Fig. 8A**). However, at Day 7, *P. bivia* burdens in cervical and uterine tissues, but not vaginal tissues, were significantly lower in ^HMb^mice compared to conventional mice (**Fig. 8B**). In comparison, no differences in vaginal swab (**Fig. 8C**) or tissue (**Fig. 8D**) burdens were observed between ^HMb^mice or conventional mice vaginally inoculated with UPEC.

**Figure 8.**
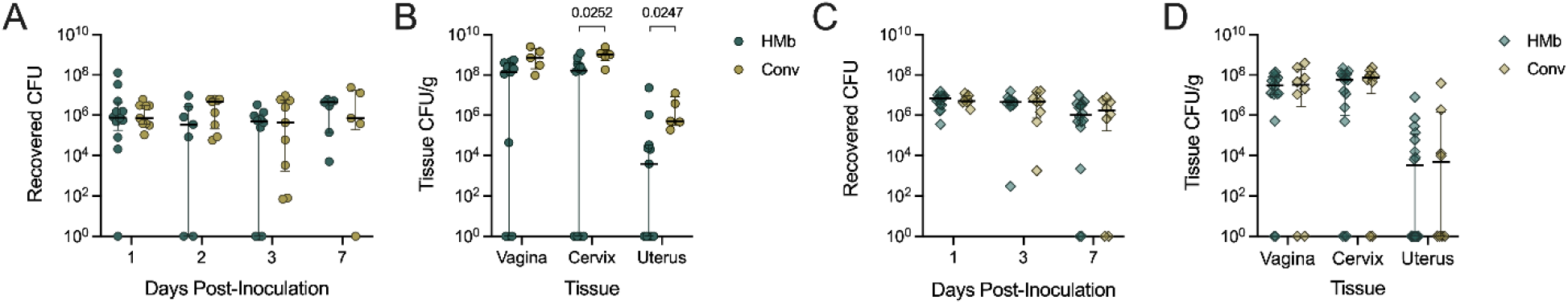
^HMb^mice have reduced cervical and uterine tissue burdens of *P. bivia* but lack protection against colonization or ascension of UPEC. ^HMb^mice and conventional (conv) mice were inoculated with 10^6^ CFU of *P. bivia* or 10^7^ CFU of UPEC. Recovered *P. bivia* CFU from **(A)** vaginal swabs collected on Days 1, 2, 3, and 7 post-inoculation and **(B)** reproductive tract tissues harvested on Day 7 (*n* = 5-12). Recovered UPEC CFU from **(C)** vaginal swabs collected on Days 1, 3, and 7 post-inoculation and **(D)** reproductive tract tissues harvested on Day 7 (*n* = 8-16). Symbols represent individual mice. Data were statistically analyzed by Mann-Whitney test and statistically significant *P* values are reported.

## DISCUSSION

Despite strong clinical correlations between the vaginal microbiota and women’s health outcomes, the ability to ascribe function of the microbiota in vaginal physiology, immunity, and susceptibility to disease is impaired by the lack of an animal model that recapitulates the human vaginal microbiota. *Lactobacillus* spp. dominate ~73% of human vaginal communities and comprise 70% of the total community abundance in humans (18) but are rare (<1%) of the vaginal communities in other mammals (62). Attempts to colonize animal models such as non-human primates (63) or laboratory mice (27–31) with human vaginal bacteria have failed to achieve long-term colonization. Key goals of this study were to examine the impact of host environment, microbial exposure, and estrous cycle on the vaginal microbiota on the same murine genetic background and to establish the influence of the vaginal microbiota on pathogen introduction in mice exposed to human microbes.

To our knowledge, this is the first comparison of the vaginal microbiota across mice of the same genetic background within and between vivaria. Similar to murine fecal communities (43, 64), we observed a strong influence of vivaria on vaginal microbiota composition in conventional C57BL/6J mice across three distinct facilities (**Fig. 1**). Moreover, we discovered high variability between vaginal compositions of mice within the same colony and effects of moving mice between rooms. Factors that may contribute to the microbial variation are likely due to a culmination of small differences in housing maintenance, diet, and bedding between facilities, or whether mice are differentially exposed to stressors such as noise, frequency of handling or transportation, or exposure to light (43, 64, 65). Congruent with our study, distinct changes to the vaginal microbiota have also been reported in wild field mice upon captivity (66).

The vaginal microbiota of ^HMb^mice, an existing humanized gut-microbiota murine model, recapitulated the community structure seen in conventional mice characterized by low alpha diversity and dominance by a single taxa in the majority of mice (**Fig. 2**). This finding suggests that host selective pressures drive the vaginal community towards a skewed dominance of a single organism independent of microbial exposure (23–25). We observed high variability between cohorts of ^HMb^mice sampled at different times from the same colony, but this variability could still be grouped into consistent ^h^mCSTs (**Fig. 4**) suggesting continuity of dominant microbes over time. Vaginal microbial fluctuation was not explained by seasonal changes or convergence of microbial compositions as reported previously (42, 67, 68) over the two-year sampling period of this study. Compared to conventional mice, ^HMb^mice were enriched in colonization by *Lactobacillus* spp.; however, there remain several limitations to this model. There is likely some “conventionalization” of ^HMb^mice since we detected a *S. succinus*-dominant community (^h^mCST II) which is the most common community present in conventional C57BL/6J Jackson mice (23, 24). Additionally, ^HMb^mice were frequently colonized by *Lactobacillus-*dominant communities (^h^mCST I and ^h^mCST V) that had OTUs mapping to *L. murinus*. While not a human-associated *Lactobacillus* sp., *L. murinus* has been isolated from the vaginal tract of wild mice (66) and from the gut of conventional C57BL/6J mice (69). Lastly, another frequently observed ^HMb^mice community was dominated by *Enterobacteriaceae* including *E. coli*(^h^mCST III). Although *E. coli* are reported in human vaginal samples, they are typically at low relative abundance in 12-27% of non-pregnant women and 14-33% of pregnant women (70–73).

In humans, the vaginal microbiota composition fluctuates over the menstrual cycle likely due to steroid hormone-mediated changes in glycogen availability, a key nutrient source for Lactobacilli and other microbes (62, 74, 75). Observations in other mammals are mixed; reproductive cycle-associated fluctuations have been reported in some studies of non-human primates, bovine species, and rats (76–80) but not in other non-human primates, horses, or mini-pigs (81–84). Like conventional C57BL/6J mice (24), we did not observe a strong influence of estrous stage on vaginal microbial compositions or individual taxa in ^HMb^mice. Counter to human studies (51), we observed a modest, but significant, fluctuation in Shannon diversity index over the estrous cycle with the lowest diversity occurring during estrus (**Fig. 3F**). An important caveat to consider is that we only qualitatively determined estrous stage based on cytology of vaginal washes and did not measure hormone levels. It is also possible that this study was underpowered to delineate distinct microbial signatures. Still, our findings are consistent with previous studies in conventional C57BL/6 mice demonstrating minimal impacts of estrous stage on the fecal (85) and vaginal (24) microbiomes.

Because the vaginal microbiota is believed to play an important role in protection against pathogens (86), we tested the impact of the altered vaginal microbiota of ^HMb^mice on vaginal colonization by GBS, a leading cause of neonatal invasive disease and agent of aerobic vaginitis (58). GBS colonization is correlated with specific taxa including *Staphylococcus* spp., *P. bivia*, and *E. coli* in non-pregnant women (72, 87), and GBS uterine ascension is correlated with *Staphylococcus*-dominant vaginal microbiota in conventional mice (24). GBS uterine ascension is a mechanism for pregnancy complications including chorioamnionitis, preterm birth, or stillbirth (88–90). Although there were minimal differences in GBS vaginal burdens between ^HMb^mice and conventional mice, ^HMb^mice consistently demonstrated lower uterine burdens and revealed six different vaginal taxa that were inversely correlated with detection of uterine GBS (**Fig. 5**). *In vitro, L. murinus*, but not *E. coli*, reduced GBS growth in coculture experiments, while exogenous treatment of *E. coli*, but not *L. murinus*, reduced GBS colonization in ^HMb^mice *in vivo* (**Fig. 6–7**). The discordance between *in vivo* and *in vitro* findings could be explained by attenuation of *L. murinus* anti-GBS activity *in vivo* due to poor colonization or insufficient production or ineffective local concentrations of anti-GBS factors. Concurrently, the *in vivo* success of exogenous *E. coli* could be explained through outcompeting GBS for key nutrients or attachment to host surfaces or the elicitation of an altered immune response. These studies highlight the complexity of host and microbial factors dictating GBS colonization success and may explain why some probiotics with potent anti-GBS activity *in vitro* have failed to reduce GBS vaginal colonization in clinical trials (91–94).

^HMb^mice were not consistently protected against vaginal pathobionts compared to conventional mice; ^HMb^mice displayed reduced cervical and uterine burdens of *P. bivia*,but not UPEC (**Fig. 8**). *P. bivia* uterine burdens in conventional C57BL/6J mice (10^4^-10^5^ CFU/g) were comparable to previous studies (95) suggesting that ^HMb^mice may actively suppress *P. bivia* uterine ascension or persistence. No differences in UPEC colonization were seen between ^HMb^mice and conventional mice; tissue burdens were consistent with previous findings (60, 96, 97). Counter to other murine gut colonization models (98), exogenous UPEC did not appear to be negatively impacted by the frequent endogenous vaginal *Enterobacteriaceae* or *E. coli* found in ^HMb^mice; however, we did not assess changes to the vaginal microbiome following UPEC inoculation, so it is unknown whether UPEC had any impact on endogenous *Enterobacteriaceae* including *E. coli*. GBS and UPEC, but not *P. bivia*, induce vaginal immune responses in conventional murine models (96, 99, 100), and an important limitation of our study is that we did not assess host immune responses to these pathobionts.

Our results reveal the plasticity of the mouse vaginal microbiota in response to environmental exposures, perhaps a more potent driver of variability than host genetics or biological factors such as estrus. Even so, open questions remain regarding the biologic factors driving rapid changes to the vaginal microbiota in mice. Although not an exact representation of the human vaginal microbiota, the ^HMb^mouse model described here is enriched in *Lactobacillus*-dominant communities and demonstrates the importance of the vaginal microbiota in shaping outcomes of reproductive tract infections. Continued improvement of humanized mouse models will provide a pathway to establish the functional role of the vaginal microbiota in health and disease and serve as an improved preclinical model for microbe-based therapies.

## METHODS

### Bacterial strains

GBS strain COH1 (ATCC BAA-1176) was grown in Todd-Hewitt Broth (THB) for at least 16 h at 37°C. Overnight cultures were diluted 1:10 in fresh THB and incubated at 37°C until mid-log phase (OD_600nm_=0.4). A spontaneous streptomycin-resistant mutant of UPEC strain UTI89 (101) was generated by plating an overnight culture on Luria Broth (LB) agar containing 1000 μg/mL Streptomycin. UPEC Strep^R^ was grown overnight in LB with 1000 μg/mL Streptomycin and washed twice with PBS prior to inoculation. *Prevotella bivia* Strep^R^ (100) was grown anaerobically (<100 ppm oxygen) in a Coy anaerobic chamber maintained at 37°C. *P. bivia* was cultured in Tryptic Soy Broth (TSB) with 5% laked, defibrinated sheep blood for three days. *E. coli* and *L. murinus* were isolated from ^Hmb^mice vaginal swabs plated on MRS agar. Prior to inoculation, *E. coli* and *L. murinus* were grown anaerobically in MRS overnight or over two days, respectively.

### Animals

Animal experiments were approved by the Baylor College of Medicine and University of California San Diego Institutional Animal Care and Use Committees and conducted under accepted veterinary standards. Mice were allowed to eat and drink *ad libitum*. Humanized Microbiota mice (^HMb^mice) were generated and maintained as described previously (34). WT C57BL/6J female mice (#000664) were purchased directly from Jackson Labs or from C57BL/6J stocks bred at BCM and UCSD. Prior to bacterial infections, mice were acclimated for one week in the biohazard room. Mice ranged in age from 2-6 months.

### Sample collection and estrous stage assignment

Vaginal swabs for 16S sequencing and estrous staging were conducted as described previously (57). Wet mounts of vaginal swab samples were observed under brightfield 100X magnification on an Echo Revolve microscope. Estrous stages were delineated by three independent researchers according to parameters described previously (102, 103) and assigned with a consensus of at least two researchers. Mice were sampled at a single time point (*n*= 2), every three days (*n*= 32), or daily (*n*= 5) over the span of seven days.

### DNA extraction and 16S rRNA V4 amplicon sequencing

DNA from vaginal swabs were extracted using the Quick-DNA Fungal/Bacterial Microprep Kit protocol (Zymo Research) and following manufacturer’s instructions with two deviations: samples were homogenized for 15 minutes during lysis, and DNA was eluted in 20 μL of water. Amplification and sequencing of the V4 region of the 16S rRNA gene were carried out by BCM Center for Metagenomics and Microbiome Research or UCSD Institute for Genomic Medicine using the Illumina 16Sv4 and Illumina MiSeq v2 2×250bp protocols as described (23, 24). Sequences were joined, trimmed to 150-bp reads, and denoised using Deblur through the QIIME2 pipeline using version 2022.2 (104). Operational Taxonomic Units (OTUs) were assigned using the naïve Bayes sklearn classier trained on the 515F/806R region of Greengenes 13_8 with 99% similarity (105).

Vaginal samples of Jackson Labs mice from our previous work (24) were downloaded from EBI accession number PRJEB25733 and included in our present study (EBI accession number PRJEB58804) for **Fig. 1** and **Fig. S2**. Since many of the samples were low biomass, DNA contaminants from sequencing reagents and kits had a substantial impact on the dataset and necessitated filtering of Feature IDs as presented in **Fig. S4**. First, feature IDs that appeared in less than seven samples were removed. Second, negative controls that went through the entire pipeline, from DNA extraction to sequencing, were run through the R package Decontam (106) (R version 4.2.0 (2022–04-22) – “Vigorous Calisthenics”), which identified 35 Feature IDs that were subsequently removed from the feature table. Lastly, the feature table was re-imported into Qiime2 where other abundant contaminants (*Streptophyta, Geobacillus, Thermus, Phyllobacteriaceae, Bradyrhizobium*, and *P. veronii*) were filtered out.

Alpha diversity (OTUs and Shannon), beta diversity (Bray-Curtis distance and PERMANOVA), and differential abundance (ANCOM) tests were carried out in QIIME2 (107). Output files were exported and analyzed in R Studio Version 1.2.5001 using factoextra (108), and Phyloseq (109). To assign ^h^mCSTs and create heatmaps, hierarchical clustering was performed using the R package stats on the filtered feature table with Ward’s linkage of Euclidean distances (23). Data visualization was performed with ggplot2 (110) and GraphPad Prism v9.4.0 (GraphPad Software, Inc., La Jolla, CA).

### Reproductive parameters and data

Data for Jackson Labs was found in the Handbook on Genetically Standardized JAX Mice (50). Data for BCM C57BL/6J and ^HMb^mice colonies were sourced from colony managers. Ranges were not provided by other vivaria but could be determined from the ^HMb^mice breeding data.

### Murine pathogen colonization models

Vaginal colonization studies were conducted as described previously (57, 95). For GBS and UPEC colonization experiments, mice were synchronized with 0.5mg β-estradiol administered intraperitoneally 24h prior to inoculation (**Fig. 5A**). For *P. bivia* colonization experiments, mice received β-estradiol 48h and 24h prior to inoculation. Mice were vaginally inoculated with 10μL of GBS COH1 (10^7^ CFU), UPEC Strep^R^ (10^7^ CFU) or *P. bivia* Strep^R^ (10^6^ CFU). Vaginal swabs were collected at indicated timepoints and tissues were harvested on day 7 as previously described (23). CHROMagar StrepB Select (DRG International Inc.) agar plates were used to quantify recovered GBS (identified as pink/mauve colonies). CHROMagar Orientation plates were used to quantify recovered UPEC (identified as pink colonies). *P. bivia* was quantified on blood agar containing 1000mg/mL Streptomycin. For pre-treatment experiments, 0.5 mg β-estradiol was given on Day −5 and −2 and either *L. murinus* (10^6^ CFU), *E. coli* (10^6^ CFU), or MRS media were administered on Day −4 and −1 before GBS challenge (**Fig. 7A**). Swabs and tissues were collected as stated above.

### *In vitro* competition assays

GBS and murine isolates of *L. murinus* and *E. coli* grown anaerobically in MRS media as cocultures or monocultures at the indicated concentrations. Samples collected at 3 and 18 hours were plated aerobically on THB (*L. murinus* competition) or CHROMagar Orientation (*E. coli* competition) plates cultured at 37°C in an anaerobic chamber for two days.

### Data Availability

Sequencing Data used in this study is available in EBI under accession number PRJEB58804. Scripts are accessible at GitHub under project “MouseVaginalMicrobiota-HMb_filtering_CST”.

### Statistics

All data were collected from at least two independent experiments unless otherwise stated. Mean values from independent experiment replicates, or biological replicates, are represented by medians with interquartile ranges or box-and-whisker plots with Tukey’s as indicated in the figure legends. Independent vaginal swab microbial communities, some taken at multiple timepoints from the same mouse, are represented by each symbol on PCoA plots. Pathogen burdens between conventional and ^HMb^mice were assessed by Mann-Whitney test. Alpha and beta-diversity metrics, and GBS burdens by ^h^mCST were analyzed by Kruskal-Wallis with Dunn’s multiple comparisons test. Competitive indexes were statistically analyzed by one sample t test with a theoretical mean of 1.0. Coculture and monoculture comparisons were performed using two-way ANOVA with Šídák’s multiple comparisons test. ^h^mCST frequencies across estrous stages were compared by Chi square test. Statistical analyses were performed using GraphPad Prism, version 9.4.0. *P* values < 0.05 were considered statistically significant.

## AUTHOR CONTRIBUTIONS

KAP and MAM conceived and designed experiments. MAM, VME, JJZ, SO, KR, and MBB performed experiments and collected data. KAP and MEM analyzed and interpreted results. MEM, VME, and KAP drafted the manuscript. KAP and RAB secured funding. All authors contributed the discussion/manuscript edits.

## ACKNOWLEDGEMENTS

We would like to acknowledge Colleen Brand of the Britton lab for colony management of the ^HMb^mice, Dr. Stephanie Fowler for providing breeding data for the BCM C57BL/6J colony, the Center for Metagenomics and Microbiology Research for 16S sequencing, and Amanda Lewis and Nicole Gilbert for the strain of streptomycin resistant *Prevotella bivia*. MEM and JZ were supported by an NIH T32 award (T32GM136554) and MEM, VME, and SO were supported by NIH F31 awards (AI167538, AI167547, and HD111236) respectively. Studies were supported by the Burroughs Wellcome Fund Next Gen Pregnancy Initiative (NGP10103), and NIH R01 (DK128053) and U19 (AI157981) to KAP. The funders had no role in study design, data collection and interpretation, or the decision to submit the work for publication.

