## Supplemental Material for "Vaginal microbial dynamics and pathogen colonization in a humanized microbiota mouse model"

### SUPPLEMENTAL MATERIALS

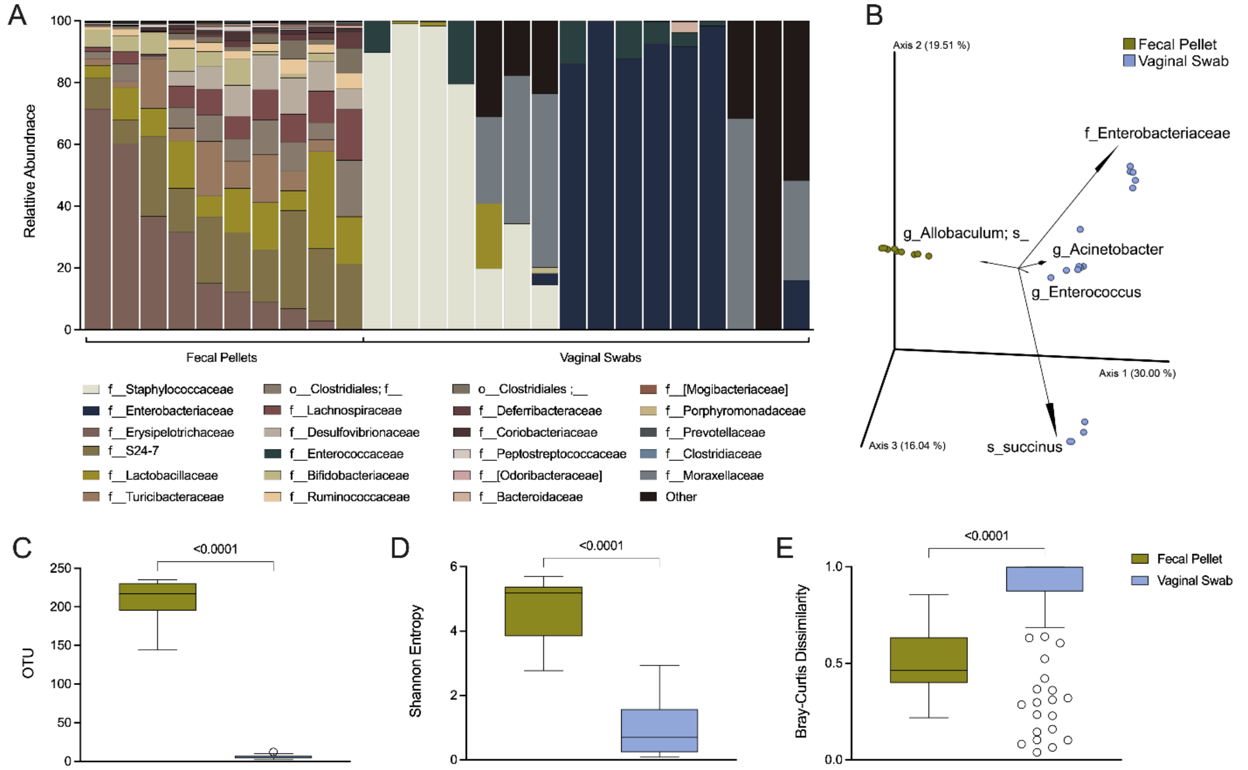


**Supplemental Figure 1. $^{HMb}$mice have tissue-specific microbial compositions.** Vaginal swab and fecal pellet sets were collected from a cohort of $^{HMb}$mice (*n*=16) **(A)** Microbial compositions of fecal pellets (left) and vaginal swabs (right). **(B)** Clustering of fecal and vaginal communities based on Bray-Curtis distances. **(C)** Observed OTUs, **(D)** Shannon Entropy, and **(E)** pairwise Bray-Curtis distances from both sample types. Symbols represent microbial profiles from individual mice (B-D) or pairwise Bray-Curtis distances generated using PERMANOVA (E) with Tukey's boxplots displayed. Data in C-E were statistically analyzed by Kruskal-Wallis with Dunn's multiple comparisons test and statistically significant *P* values are reported.

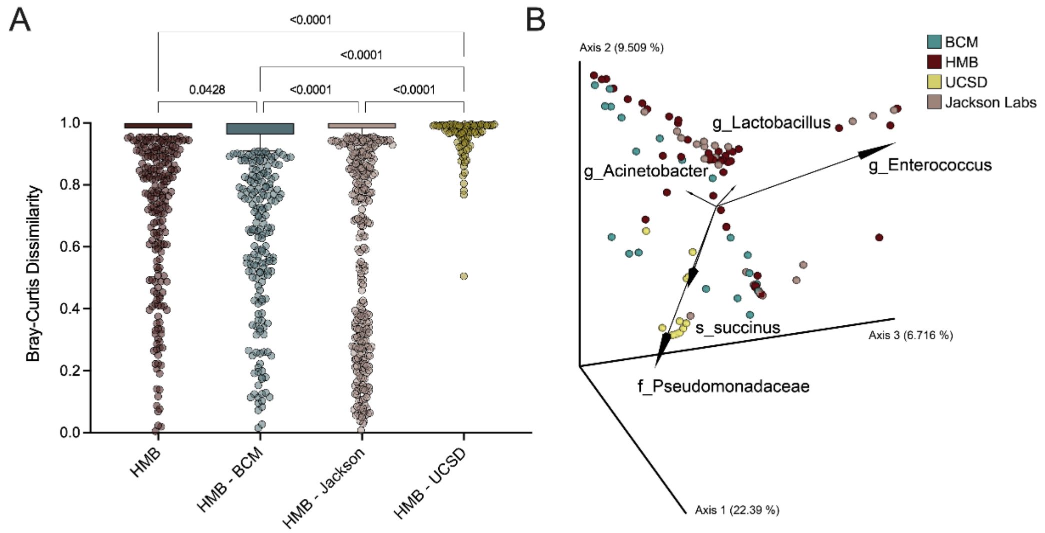


**Supplemental Figure 2. HMbmice display unique and variable vaginal microbiota compared to conventional mice with some shared taxa including *Lactobacillus*, *Enterococcus*, and *Staphylococcus*.** HMbmice vaginal compositions were compared to the vaginal microbiota of the conventionally colonized mice from **Figure 1**. **(A)** Intra-site dissimilarity based on Bray-Curtis distances. **(B)** Clustering of vaginal communities from mice in each colony based on Bray-Curtis distances. Symbols **(A)** represent pairwise comparisons or **(B)** vaginal communities from individual mice. Data were statistically analyzed with Kruskal-Wallis and Dunn's multiple comparisons test and statistically significant *P* values are reported.

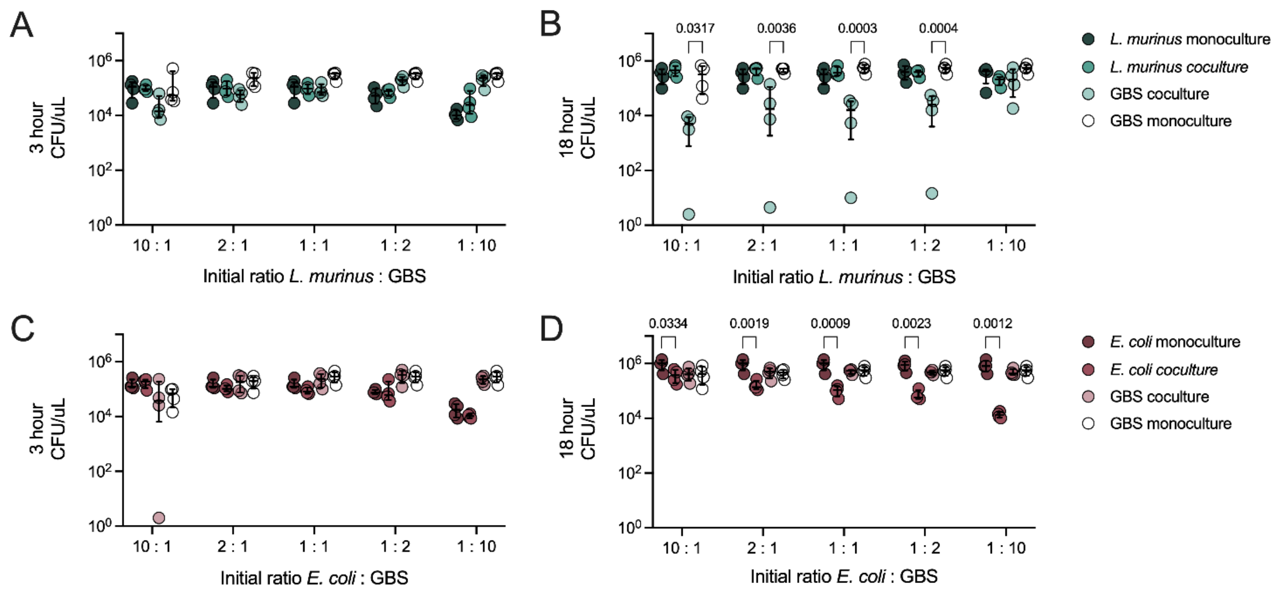


**Supplemental Figure 3. Viable CFU recovered from GBS competition experiments with *L. murinus* and *E. coli*.** GBS was cocultured with either **(A, B)** *L. murinus* or **(C, D)** *E. coli* at increasing concentrations of GBS. Viable CFU of all microbes in coculture as well as monoculture controls were quantified at **(A, E)** 3 hours and **(B, D)** 18 hours. Comparisons between monoculture and coculture were analyzed by 2-way ANOVA with Šídák's multiple comparison test; statistically significant *P* values are shown.

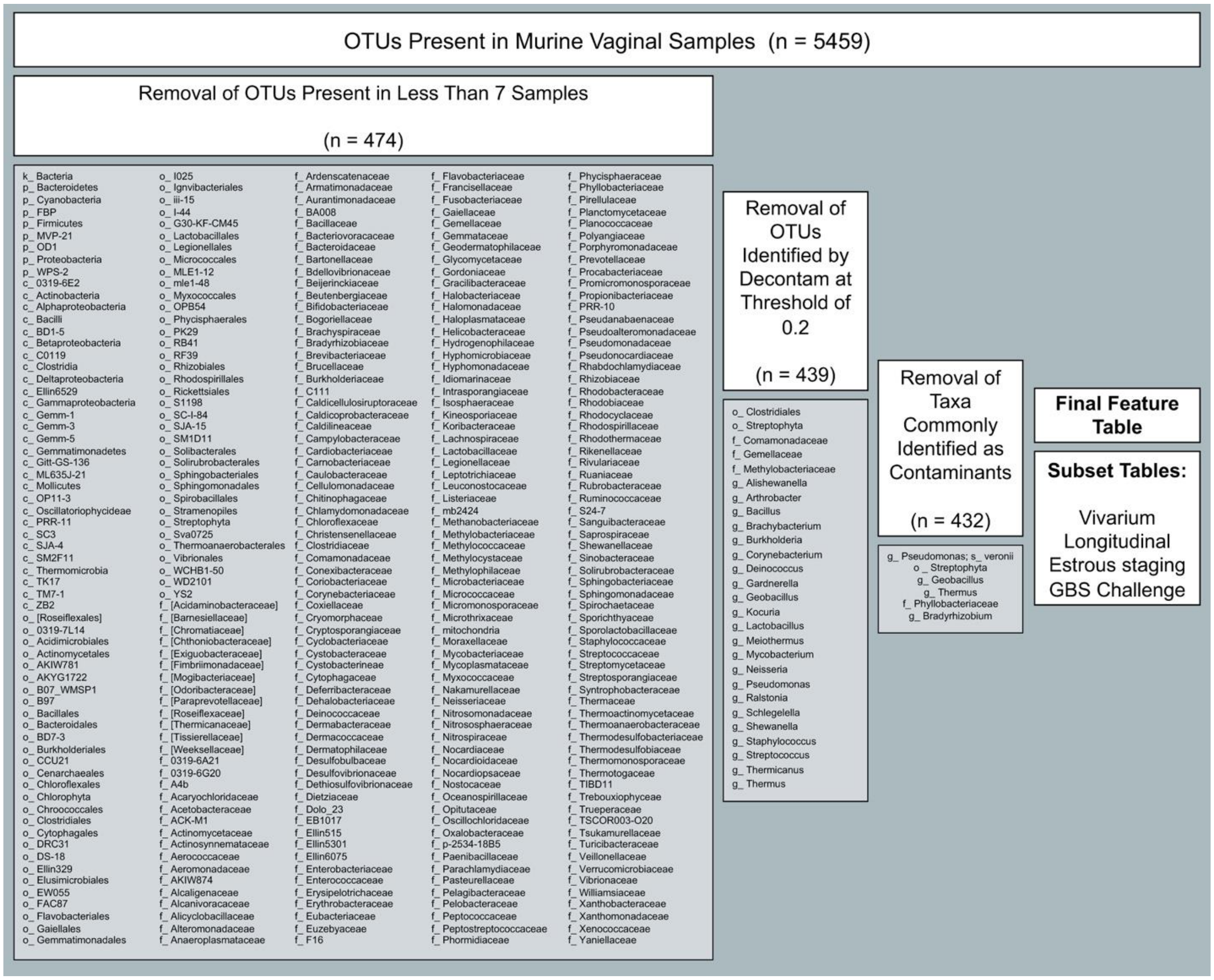


**Supplemental Figure 4. Flow chart of contaminant sequence removal to generate feature tables.** All studies were merged prior generation of the taxonomy list. In the resulting feature table, OTU removal was performed first: OTUs that appeared in less than 7 samples were removed in Qiime2 followed by the removal of contaminants using Decontam in R. The table was re-imported into Qiime2 for the manual removal of known taxonomic calls identified from prior studies. The final feature table was then subset accordingly.

**Supplemental Table 1. *Lactobacillus* species candidates for** $^{h}$mCST-associated **OTUs cross-referenced through BLAST[a].**

| Taxonomy | OTU | BLAST result[b] | $^{h}$mCST |
|---|---|---|---|
| g_Lactobacillus | 036cd775b052774d0d142aecfff4fef3 | L. fermentum | I |
| | 1751fd89760ef9bcde0c89ea1ec7dad7 | | |
| | 28017a3bb16d6e95eab4f16d1102872a | | |
| | 95854254d37f27a4369d20d3ff9392ca | | |
| | 9938c8f22908861f780f8719b75dc0b2 | | |
| | 9e5e21ee5e1909019da5f5fae2cf9def | | |
| | be2298b15183a60c07593ffa57c74eb3 | | |
| | f2b4d483605d38c455115c7aeb704a3b | | |
| | f5d5bf98f708c7d8e30d086a07a99265 | L. fermentum<br>L. mulieris<br>L. delbruekii<br>L. femina | |
| | 03cc6d9b865904a295617ca4aaa4b24f | L. gasseri<br>L. paragasseri | |
| | 1b5257c442e29ac8b820cfd82d00f1cd | | |
| | 249492182aa51dfbc16ef97b9c46f4c8 | L. gasseri<br>L. paragasseri<br>L. johnsonii | |
| | 2a76b69853ab0b8a89e17694e7a6ebff | | |
| | 50f60ff0d1f3211f4b43eebbeeb07cf4 | | |
| | 525e1879094cddd4a7b36239088ff0ea | | |
| | 5a5fa98df77b3a334e9d0eb2ffabd26c | | |
| | 7ed1d6eee61436b8aa293945a4809b1b | | |
| | 939c9bb217ad250af43b6dd6fed184fb | | |
| | 96f31c5559dba8d4ff9b58376d59254c | | |
| | b3fef1e26b7b6ac20847ede4da68547a | | |
| | bdb0c76ab13a64c1d0abc4606f4ee255 | | |
| | d1cd930a42c11eab0f3a3f7a875b5b07 | | |
| | e4e5a0667e88f7617f4875e80d790bb5 | | |
| | 1fb2f4a7f74d291e5f8d7d5b20790732 | L. jensenii | |
| | ba70ebcf8e200d42b40dd856850d6440 | | |
| | 7583e1d29147d3cbc473a2289aa19787 | L. johnsonii<br>L. gasseri<br>L. paragasseri | |
| | eac1498be3c53337b2586b49bcf309a9 | | |
| | 47cc8d7e1ab2c127c73fecb5767dc7b9 | L. kimchicus<br>L. apodemi<br>L. brevis | |
| | 34f5e40f9fb2818d0692b993e5c9b923 | L. murinus | |
| | 87a9b78f342b92b99ed153d2256845f6 | | |
| | 34e8216d377ee04e2841dec21f857a88 | Uncultured<br>L. fermentum | |
| | 947b0003a4e55351b91765a8a1900586 | Uncultured<br>L. gasseri | |
| | a515079065dc5c33f95af12b743886c3 | Uncultured<br>L. gasseri | |

| | | | |
|---|---|---|---|
| | f002e38adcccc245fb0102cd84080014 | Uncultured L. intestinalis | |
| | 94ea9e9e8b1b060ea09df186e05065f1 | Uncultured L. murinus | |
| g_Lactobacillus; s_ | 388d68047722cd72012b8b00141d0fcd | L. aviarius and L. araffinosis | V |
| | f49f940159fa9e43f5e78335994e2dc6 | | |
| | 1d0ff07e34de73fbbdd96097c589578c | L. buchneri L. diolivorans L. hilgardii L. farraginis L. rapi L. sunkii | |
| | bde5a716b4942131d93365d3d8ddd769 | L. fermentum | |
| | 0c1012b02835602b17b5c366461212af | L. murinus | |
| | 62e9d424e478e73e21c41e6e61e34866 | | |
| | 97b0ff875799fa576ddf5a0f9dafc87f | | |
| | bbfade6b215de60d0f119bdd8f66286e | | |
| | d3211b7a04e5fb36a383450b8f5710e2 | | |
| | e6dfb454c58aeef460b0f37c800400d3 | | |
| | 5232064f3b3da2958610b16281db5ec4 | L. pobuzihii L. murinus L. salivarius | |
| | 27192809551816b03fc90fa4edea030e | L. reuteri | |
| | 41558c8fedbc65e50d5d5ba12da7f231 | | |
| | c348a7cc121fa43fb14ca92cc4273c2e | | |
| | db7a83e6eedefdbd44c5a2256967b705 | | |

[a]Timestamp for BLAST search is Jan. 12, 2023 at 00:54:14.

[b]Many results denoted “Uncultured bacterium” as a result.
